# Population trajectory analysis reveals divergent state-space geometries across cortical excitatory cell types

**DOI:** 10.1101/2025.08.13.670197

**Authors:** Zihan Yang, Yuchen Xiao

## Abstract

Cortical population activity unfolds along structured trajectories in state space, reflecting underlying dynamics of the neural networks involved. However, how such geometry differs across neural cell types remains poorly understood. Using a large-scale two-photon calcium imaging dataset from the Allen Brain Observatory, we characterized the population trajectories of three cortical excitatory subtypes (Cux2, Emx1, and Slc17a7). These cells constitute most of the layer 2/3 excitatory populations, span distinct projection and functional hierarchies, and collectively orchestrate cortical integration and excitatory signaling. While viewing drifting gratings, Cux2 neurons exhibited compact and stable trajectories, whereas Emx1 and Slc17a7 showed broader and more distributed dynamics, implying their functional divergence in visual processes. Such geometric distinctions can be mechanistically accounted for within a low-dimensional control space where the dominant axes correspond to the effective integration timescale and recurrent gain. This geometric–mechanistic framework provides a generalizable route to bridging cortical dynamics with fundamental, cellular identity.

## Introduction

Understanding how different cortical cell types shape the geometry of population dynamics is crucial for explaining how cellular diversity gives rise to distinct dynamical regimes within the cortex. The cortical network comprises multiple neuronal types defined by genetic markers, projection targets, and intrinsic physiological characteristics^1^. These neuronal subtypes are embedded in recurrently connected circuits that constitute an information-processing architecture both hierarchical and recursive in nature^2^. Within this architecture, stable yet tunable dynamic activity patterns emerge from the coordinated regulation of local excitation–inhibition balance^3^, cross-layer feedback, and long-range projection^4^. Differences in synaptic connection probability, short-term plasticity, and membrane time constants across subtypes confer distinct dynamical roles within population activity—some providing sustained drive or background maintenance, while others mediate rapid transitions between network states^5^. Although anatomical and molecular studies have delineated this rich structural diversity^6^, how these static features are transformed into dynamic, population-level organizations of cortical activity remains unclear. In particular, how distinct excitatory subtypes sculpt their characteristic geometric and temporal structures within cortical dynamics has not been systematically analyzed. This study thereby begins from the geometry of population activity, aiming to trace how macroscopic dynamical features emerge from and reflect physiological heterogeneity at the cellular level.

State-space analysis provides a principled framework for examining population dynamics by embedding high-dimensional neural activity into trajectories that capture coordinated temporal evolution^7^. This geometric formalism has profoundly advanced our understanding of motor generation^8^, sensory encoding^9^, and cognitive control^10^ by revealing how population activity evolves along low-dimensional manifolds—from preparation-to-execution sequences in motor cortices to task-state transitions in prefrontal circuits. Within these manifolds, trajectory curvature, divergence, and looping structure provide insights into how internal network states evolve and reconfigure over time^11^. Yet, most existing studies treat recorded neurons as a homogeneous ensemble^12^, neglecting potential cell-type-specific contributions. Such simplification risks obscuring systematic, subtype-dependent patterns in dynamic organization. Although recent work has begun to explore the population geometry of different cortical layers and regions^7,13,14^, comprehensive analyses at the level of genetically-defined excitatory subtypes remain lacking, constraining our understanding of how cellular identity determines and constrains network dynamics.

In the sensory cortex, excitatory neurons comprise multiple subtypes distributed across layers^2^. Cux2-, Emx1-, and Slc17a7-expressing neurons represent the major lineages of the cortical excitatory system. These cells exhibit diverse projection patterns and synaptic properties, functioning in neural cellular processes like excitatory differentiation and glutamatergic transmission. Cux2-expressing neurons, especially those located in upper layers, primarily form intratelencephalic projections, whereas Emx1 and Slc17a7 populations encompass broader excitatory classes spanning different laminar and projection domains^15^. Knockouts of these genes lead to circuit- and projection-specific abnormalities, underscoring their essential and nonredundant roles in cortical development and excitatory signaling^16^. These subtypes likely operate under distinct dynamical regimes^4^: upper-layer Cux2 neurons may sustain long-timescale integration and recurrent stability, whereas Emx1 and Slc17a7 populations may display faster, gain-driven dynamics that enable flexible transitions between states. However, how such subtype-dependent physiological characteristics influence population-level activities remains largely unknown.

In this study, we systematically analyzed large-scale two-photon calcium imaging data from the Allen Brain Observatory^17^ to characterize the population trajectory geometry of three cortical excitatory subtypes^18^—Cux2-CreERT2, Emx1-IRES-Cre, and Slc17a7-IRES2-Cre—located in layers 2/3 of the mouse primary visual cortex^19^. The imaging data were recorded while mice were viewing drifting gratings as visual stimuli. We constructed an integrative geometric feature framework spanning five complementary dimensions—geometric morphology, dynamic intensity, manifold orientation, coverage scope, and recursive stability—to capture the structural organization of population trajectories. This multidimensional analysis revealed that Cux2 populations exhibited compact and temporally stable trajectories, whereas Emx1 and Slc17a7 populations displayed broader, more dispersed dynamics, indicating subtype-dependent differences in state-evolution mechanisms. These distinctions remained robust across visual stimulus orientations and experimental sessions, suggesting that they arise from intrinsic biophysical and circuit-level properties rather than input-driven variability.

To mechanistically interpret these geometric divergences, we used a low-dimensional control-space model^20^ whose principal axes correspond to the effective integration timescale (τ_eff) and the effective recurrent gain (g_eff). These two parameters jointly define the dynamic regime of the network: τ_eff determines the temporal span of information integration and the smoothness of population trajectories, while g_eff modulates the amplification of recurrent feedback and the extent of trajectory expansion. By positioning each subtype within this control space^21^, we quantitatively map empirical geometric features onto underlying electrophysiological mechanisms, thereby establishing a direct link between cellular physiology and population computation. This geometry–mechanism framework reveals that excitatory subtypes occupy distinct regions of the cortical dynamical landscape and collectively form a structured continuum along the τ_eff–g_eff axis. Together, these findings provide a unified theoretical foundation for understanding how structural diversity at the cellular level gives rise to the rich repertoire of population dynamics observed across the cortex.

## Results

### Part 1 Quantitative characterization of population trajectory dynamics

#### 1.1 Trajectory geometry differs across excitatory subtypes along effective dynamical dimensions

We began by characterizing the geometric organization of excitatory population trajectories across cortical subtypes (Fig. 1A–H). Session-level trajectories were first projected into the top three principal components (PC1–PC3) of the population latent space^7^, capturing the dominant axes of temporal evolution in neural activity^22^. Representative examples revealed subtype-specific geometric envelopes: Cux2-CreERT2 trajectories remained compact and cyclic within a confined manifold, Emx1-IRES-Cre trajectories exhibited intermediate expansion and curvature, and Slc17a7-IRES2-Cre trajectories spanned the broadest regions of state space (Fig. 1A–C).

**Figure 1.**
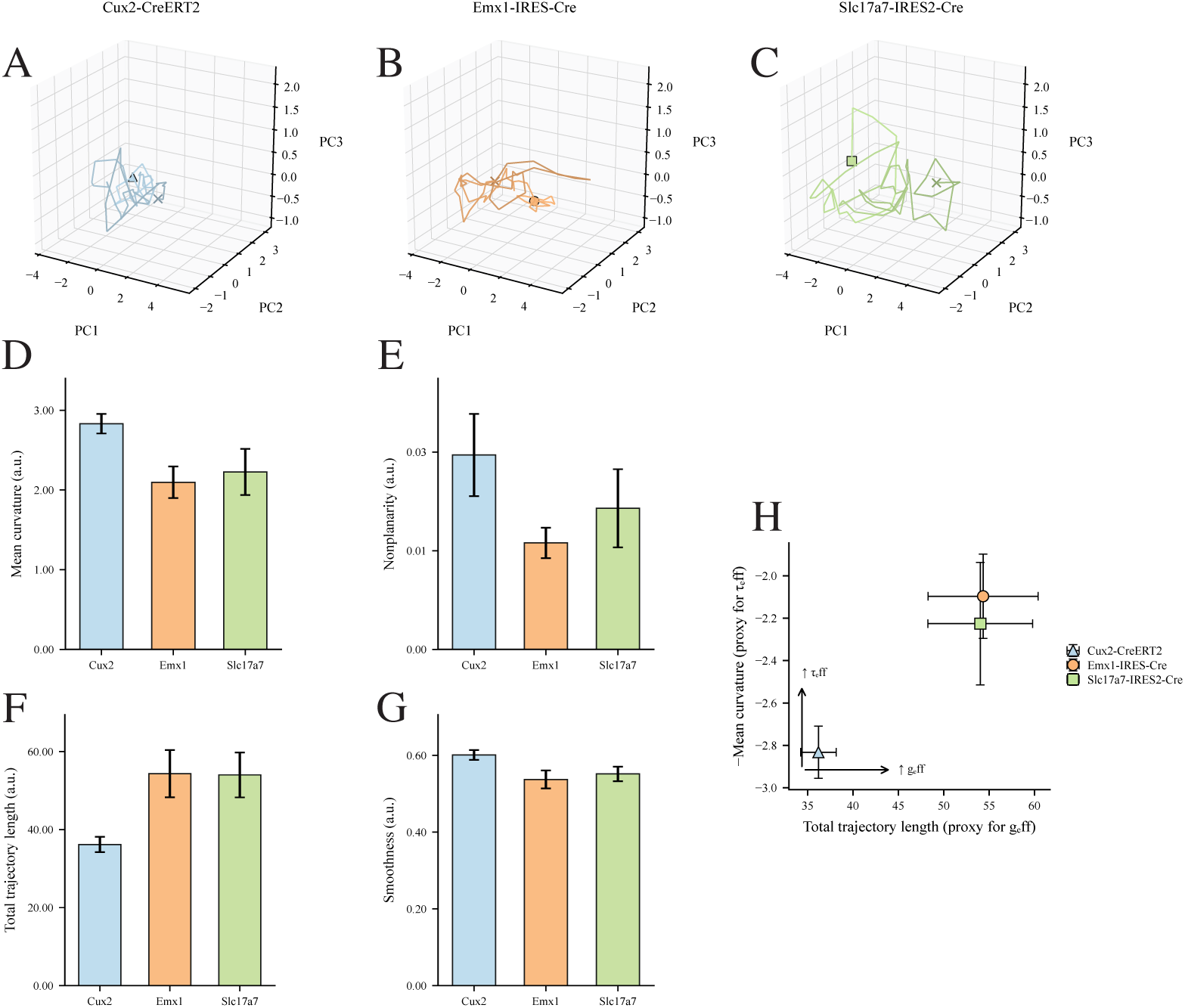
Subtype-specific geometries positioned along the τ_eff–g_eff continuum. (A–C) Representative latent trajectories (PC1–PC3) of three excitatory subtypes in mouse visual cortex. Cux2 neurons exhibit compact, cyclic trajectories; Emx1 neurons show intermediate expansion and curvature; Slc17a7 neurons display broad, dispersed dynamics. (D–G) Group-averaged geometric metrics (mean ± SEM) for curvature (reported as –κ^−^), nonplanarity (3D deviation from a local plane), total trajectory length (L_total), and short-timescale smoothness. (H) Cross-metric relationships between L_total and –κ^−^ reveal a structured gradient that aligns with the effective integration–gain (τ_eff–g_eff) parameter space, placing the three subtypes along a continuum from more stable to more flexible dynamical regimes.

To quantify these differences, we computed four structural metrics for each recording session: mean curvature as a measure of local trajectory bending, nonplanarity as a proxy for three-dimensional spread, total trajectory length (L_total) as a measure of path extent, and smoothness capturing short-timescale stability (Fig. 1D–G)^23^. Across metrics, subtypes displayed a shared hierarchical pattern, with Cux2 showing the smallest trajectory extent, Emx1 intermediate values, and Slc17a7 the largest. This pattern was robust across orientations and sessions, suggesting that the observed hierarchy reflects intrinsic circuit properties rather than stimulus-specific modulation.

Finally, we examined cross-metric relationships to map these geometric signatures onto an effective parameter space (Fig. 1H). Across sessions, trajectory length correlated inversely with mean curvature, delineating a structural continuum interpretable as a τ_eff–g_eff plane: populations with larger L_total (broader trajectories) correspond to higher effective gain (g_eff), while those with lower curvature magnitude reflect longer integration timescales (τ_eff). Within this continuum, Cux2 occupies the low-gain, long-timescale corner, whereas Emx1 and Slc17a7 lie in higher-gain, shorter-timescale regions, with Slc17a7 showing the strongest gain shift.

Together, these results establish a geometric hierarchy among excitatory subtypes, revealing that the spatial structure of population trajectories encodes an intrinsic balance between integration and recurrent amplification.

#### 1.2 Temporal and directional organization of population dynamics

We next examined how excitatory subtypes differ in their temporal and directional regulation of population activity (Figs. 2–3). Session-level trajectory derivatives were used to extract instantaneous speed (μ_v), acceleration (μ_a), and speed variance (σ_v), providing quantitative proxies for dynamic intensity and temporal stability^24^. Across subtypes, mean speed and acceleration showed a general trend of Cux2 < Emx1 and Slc17a7, with the latter two exhibiting largely comparable values (Figs. 2A–C), indicating progressively stronger activity modulation and faster temporal evolution from upper- to deeper-layer populations. Speed variance further revealed subtype-dependent stability differences, with Cux2 trajectories remaining the most stable, whereas Slc17a7 trajectories exhibited larger fluctuations—consistent with stronger recurrent drive and higher effective gain (g_eff).

**Figure 2.**
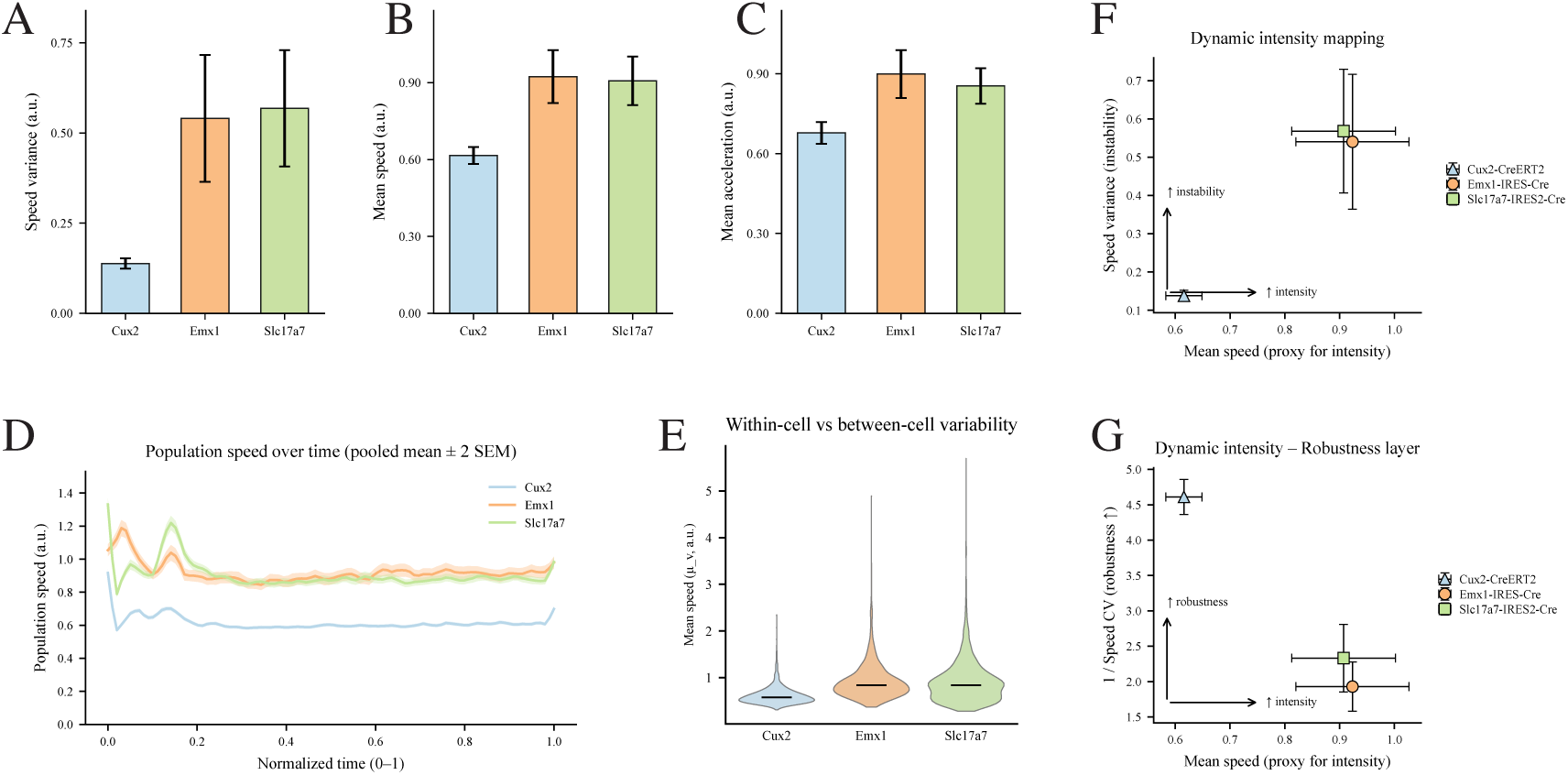
Integrative mapping of population dynamic intensity and stability across excitatory subtypes. (A–C) Group-averaged velocity (μ_v), acceleration (μ_a), and speed variance (σ_v) for the three excitatory subtypes (mean ± SEM). Cux2 populations show lowest overall values, whereas Emx1 and Slc17a7 exhibit higher dynamic intensity and fluctuation magnitude. (D) Time-course of mean velocity during stimulus presentation. Subtypes maintain stable amplitude differences throughout the stimulus period. (E) Violin plots of normalized speed distributions illustrating subtype-dependent differences in fluctuation range and distribution shape. (F–G) Cross-metric scatter relationships between dynamic intensity (velocity) and two structural measures (curvature, trajectory length). Subtypes occupy distinct regions within these relationships, reflecting consistent differences in their dynamic profiles.

**Figure 3.**
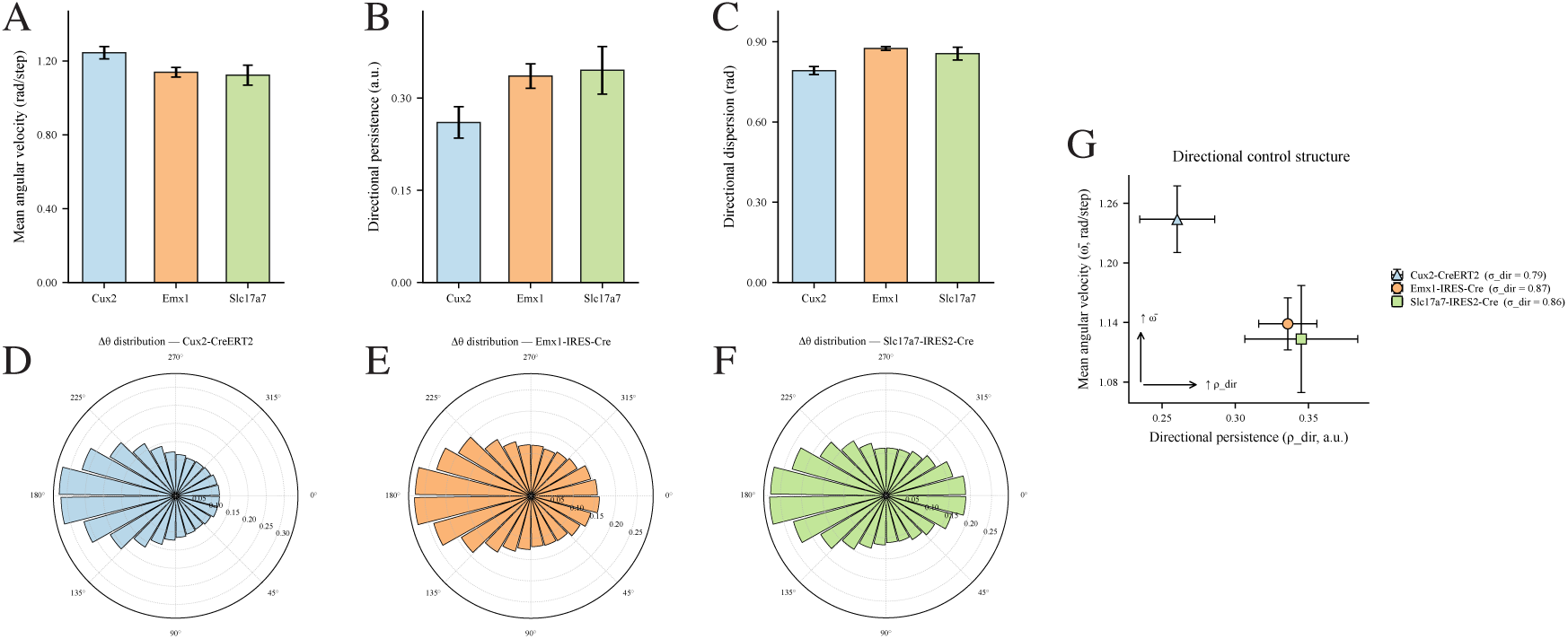
Directional control structure of population trajectories reveals subtype-specific regulation of angular dynamics. (A–C) Group-averaged directional metrics (mean ± SEM) for directional persistence (ρ_dir), mean angular velocity (ω^−^), and directional dispersion (σ_dir). Cux2 shows lower persistence and higher angular variability, Slc17a7 shows stronger directional coherence and slower reorientation, with Emx1 intermediate. (D–F) Orientation-resolved angular-displacement histograms. Cux2 trajectories span broad and rapidly shifting directions, whereas Emx1 and Slc17a7 show progressively more concentrated directional patterns. (G) Directional control plane defined by ρ_dir and ω^−^. Subtypes occupy distinct regions along a gradient from low-persistence, high-velocity dynamics (Cux2) to high-persistence, low-velocity dynamics (Slc17a7).

To integrate these measures, we mapped mean speed against speed variance, defining a dynamic control space that captures the trade-off between population intensity and stability (Fig. 2D)^25^. Within this plane, subtypes occupied distinct, non-overlapping regions: Cux2 neurons clustered near the low-intensity, high-stability corner (low g_eff, long τ_eff), while Slc17a7 neurons concentrated near the high-intensity, low-stability regime (high g_eff, short τ_eff), with Emx1 neurons positioned intermediately. Temporal evolution analyses confirmed that these intensity differences emerge early and remain consistent throughout the trial (Fig. 2E), and within-cell variability analyses (Fig. 2F) revealed broader dynamic dispersion in Slc17a7 populations.

Together, these results delineate a robustness–intensity continuum (Fig. 2G), establishing a temporal framework that links macroscopic population dynamics to effective circuit parameters.

To further probe directional organization, we quantified instantaneous angular displacements (Δθ)^11,26^ of population trajectories and derived two complementary metrics: directional persistence (ρ_dir) and mean angular velocity (ω^−^) (Fig. 3A–C). Persistence quantifies how stably a trajectory maintains its direction over time, whereas angular velocity reflects the rate of reorientation in latent space. Cux2 trajectories showed low persistence and high angular variability, while Slc17a7 trajectories exhibited sustained directional coherence and slower reorientation (Figs. 3A–F), indicating subtype-specific control over the flow of population states. When projected into a directional control plane defined by ρ_dir and ω^−^ (Fig. 3G), the three subtypes aligned along a clear gradient: populations with greater persistence and lower angular velocity corresponded to longer integration timescales (τ_eff), whereas those with higher angular velocity mapped to higher recurrent gain (g_eff).

These analyses reveal that temporal and directional control jointly structure the dynamic landscape of excitatory populations, anchoring their activity regimes within the τ_eff–g_eff continuum.

#### 1.3 Recurrence and coverage analyses reveal structured diversity in excitatory population dynamics

We further investigated the structural diversity of cortical population dynamics through recurrence and coverage analyses (Figs. 4–5). Recurrence-based measures quantify how frequently and predictably population trajectories revisit similar regions of latent space, providing an index of dynamical stability^27^. Across subtypes, recurrence rate (RR), determinism (DET), and mean recurrence time (T_rec) showed broadly comparable patterns (Figs. 4A–C): Cux2 populations displayed comparatively shorter recurrence periods, consistent with more rapid returns to similar states within a compact dynamical regime, whereas Slc17a7 populations showed comparatively longer recurrence periods and modestly altered recurrence structure. Emx1 populations again occupied an intermediate regime, suggesting graded stability along the cortical depth axis.

**Figure 4.**
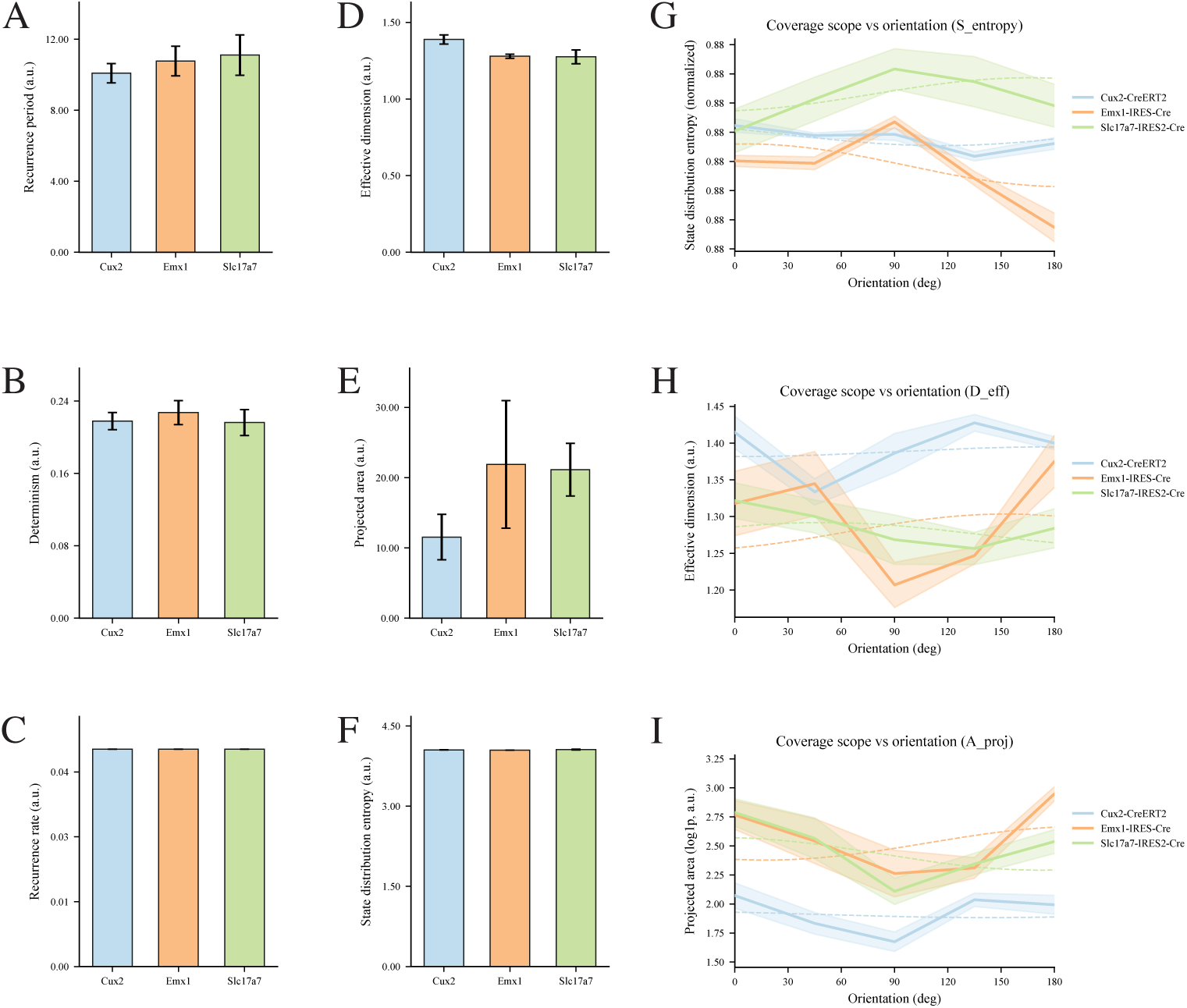
Integrative mapping of recurrence and coverage metrics reveals structured diversity in cortical population dynamics. (A–C) Session-averaged recurrence metrics for Cux2-, Emx1-, and Slc17a7-labeled populations, including recurrence rate (RR), determinism (DET), and recurrence period (T_rec). (D–F) Session-averaged coverage metrics for the same subtypes, including projected area (A_proj), effective dimension (D_eff), and state-space entropy (S_entropy). (G–I) Orientation-dependent profiles of A_proj, D_eff, and S_entropy for each subtype, showing mean values and session-wise variability across stimulus directions.

**Figure 5.**
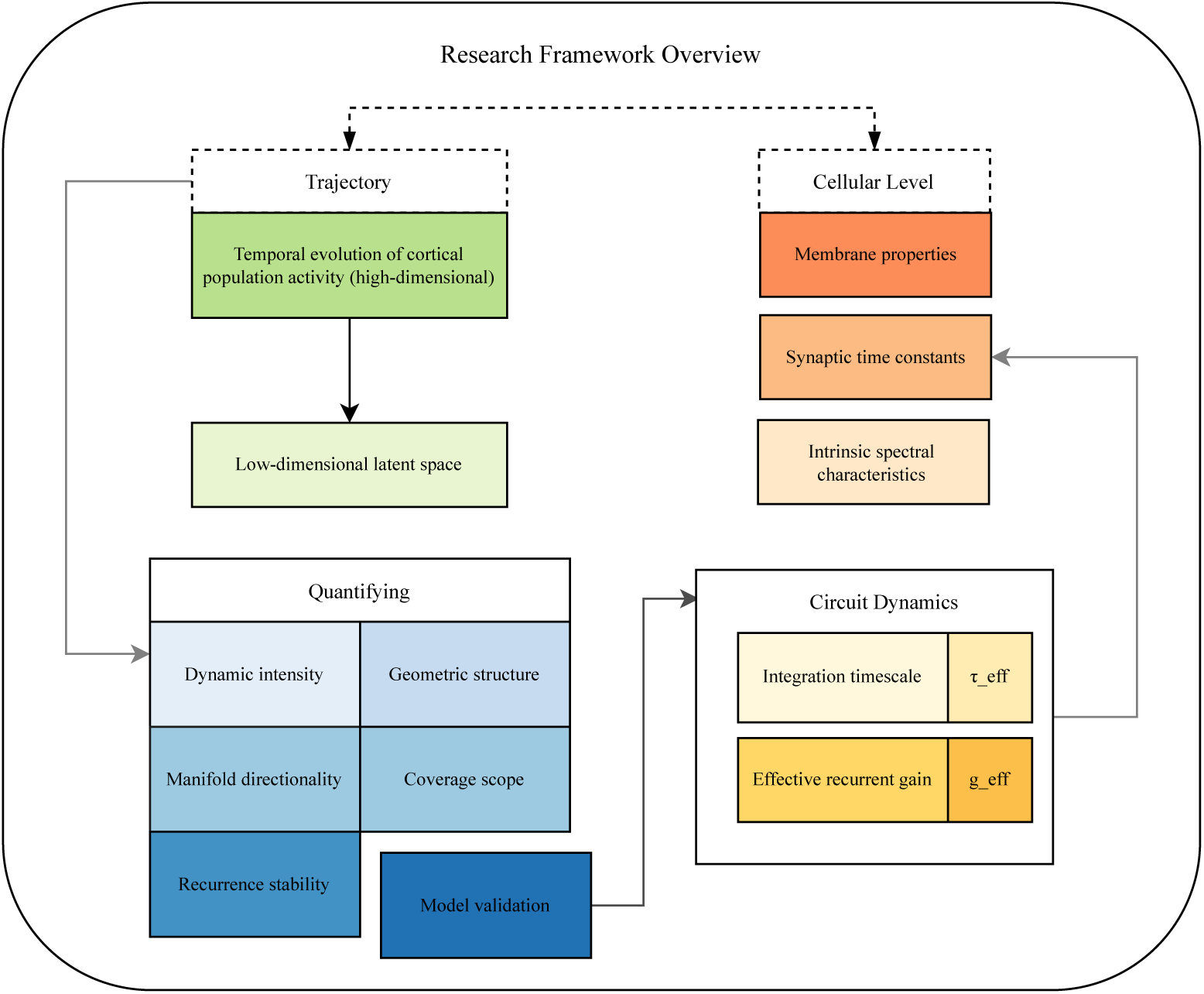
Research framework linking population trajectory geometry to circuit and cellular mechanisms. (A) Population trajectories represent the temporal evolution of high-dimensional neural activity, projected into a low-dimensional latent space for geometric characterization. (B) A set of structural metrics—dynamic intensity, geometric structure, manifold directionality, coverage scope, and recurrence stability—quantify distinct aspects of trajectory organization. (C) These metrics are mapped onto circuit-level dynamical parameters, including the effective integration timescale (τ_eff) and recurrent gain (g_eff), reflecting underlying network stability and excitability. (D) Circuit-level properties are further constrained by cellular mechanisms such as membrane and synaptic time constants and spectral properties, forming a hierarchical link from cellular to population scales.

To assess the spatial extent and diversity of state-space coverage, we quantified three complementary metrics^9^ for each session: effective dimensionality (D_eff), projected area (A_proj), and normalized state entropy (S_entropy) (Figs. 4D–F). These measures capture, respectively, the degrees of freedom engaged by the trajectory, its two-dimensional spatial footprint, and the dispersion of occupancy across the latent manifold. Cux2 trajectories were characterized by lower dimensionality and entropy, whereas Slc17a7 trajectories exhibited higher values for all coverage metrics, reflecting a broader sampling of latent states and greater representational diversity. Together, these results indicate that cortical excitatory subtypes operate within distinct dynamical regimes, ranging from compact, recurrent attractors (Cux2) to expansive, exploratory trajectories (Slc17a7).

Orientation-resolved analyses further revealed that these coverage properties are modulated by stimulus geometry (Figs. 4G–I). Entropy and dimensionality profiles varied systematically with orientation (0–180°), with Cux2 populations maintaining constrained coverage across orientations, while Emx1 and Slc17a7 populations exhibited broader and more orientation-dependent modulation. This orientation-dependent modulation indicates that population coverage diversity not only reflects intrinsic circuit properties but also interacts with feature-specific inputs.

Together, these results reveal that excitatory subtypes form a structured continuum of representational flexibility: upper-layer populations maintain stable, low-diversity manifolds, while deeper-layer populations explore broader state spaces supporting adaptive encoding.

Finally, these empirical metrics were integrated into a multilevel research framework linking population geometry to circuit and cellular mechanisms (Fig. 5). Population trajectories represent the temporal evolution of high-dimensional neural activity projected into a low-dimensional latent space, within which five structural metrics—dynamic intensity, geometric structure, manifold directionality, coverage scope, and recurrence stability—quantify distinct dynamical facets. These structural descriptors map onto circuit-level parameters, particularly the effective integration timescale (τ_eff) and recurrent gain (g_eff), which define the balance between integration and excitation in local networks. At a deeper level, these circuit properties arise from cellular determinants such as membrane and synaptic time constants and spectral characteristics, forming a hierarchical linkage from cellular to population scales that constrains how cortical circuits generate and maintain structured dynamics.

### Part II. Mechanistic analysis of the effective timescale–gain model

#### 2.1 Parameter–metric mapping and dynamical robustness analysis

To mechanistically validate the empirical τ_eff–g_eff mapping derived from population geometry, we implemented a minimal recurrent dynamical model that explicitly parameterizes integration timescale (τ_eff), recurrent gain (g_eff), internal noise amplitude (σ_int), and a plasticity-related weight modulation factor (Δw) (Fig. 6)^25^. For each parameter, systematic sweeps were performed across a ±50% range around baseline values, and three representative structural metrics—trajectory length (L), mean speed (μ_v), and mean curvature (reported with sign flipped)^28^—were computed from the simulated trajectories (16 random network realizations per condition).

**Figure 6.**
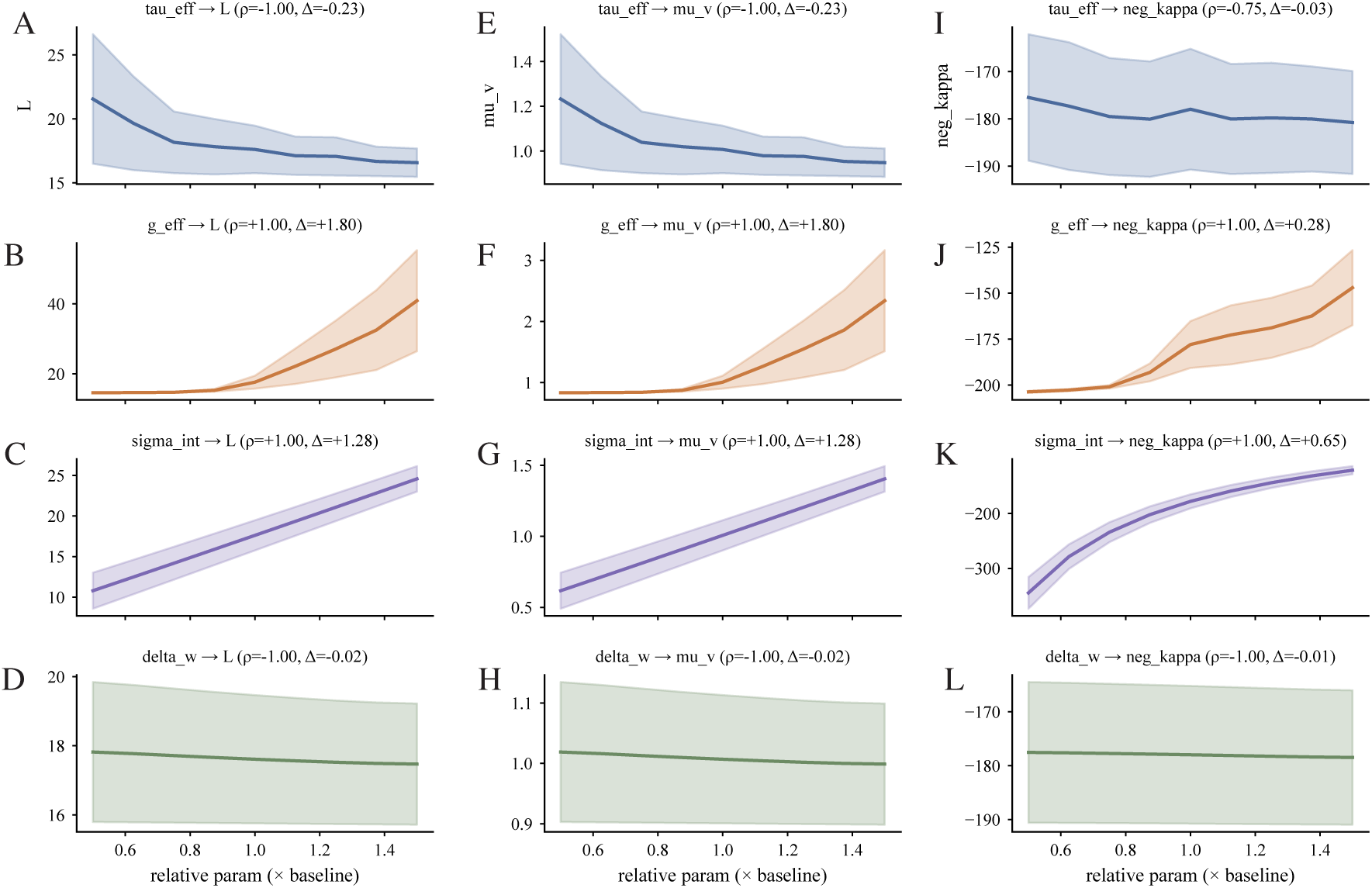
Mechanistic validation of the τ_eff–g_eff framework reveals consistent parameter–metric alignment. (A–L) Systematic parameter sweeps demonstrate monotonic relationships between mechanistic parameters (τ_eff, g_eff, σ_int, and Δw) and key structural metrics (trajectory length L, mean speed μ_v, and mean curvature (reported with sign flipped)). Each row represents one parameter, and each column corresponds to a different structural metric. Curves show mean ± SEM across 16 random network realizations. Spearman’s ρ and relative change (Δ) values denote the directionality and magnitude of parameter–metric coupling.

Each parameter exhibited a consistent and interpretable relationship with the structural metrics (Figs. 6A–L). Increasing τ_eff reduced both trajectory length and mean speed while slightly decreasing curvature magnitude, indicating slower evolution and smoother, more integrated dynamics. In contrast, increasing g_eff produced longer and faster trajectories with greater curvature magnitude, consistent with enhanced recurrent amplification and reduced integration stability. σ_int primarily modulated curvature and variability, broadening the dynamical range without strongly altering the mean trajectory length. Δw had minimal influence under the current regime, indicating that low-rank modulatory perturbations contribute weakly to the steady-state geometry. Across all parameter–metric pairs, the direction of change and Spearman’s ρ values confirmed monotonic alignment between model parameters and their corresponding empirical proxies—L and μ_v for g_eff, and –κ for τ_eff—thus reproducing the structure observed in experimental data.

These results establish a direct mechanistic correspondence between the effective circuit parameters and geometric descriptors of population trajectories. The model demonstrates that τ_eff and g_eff jointly define a low-dimensional control space governing both the speed and stability of neural population evolution, while σ_int and Δw act as modulators of stochastic variability and plastic adaptation. This quantitative parameter–metric alignment provides the foundation for the subsequent robustness and sensitivity analyses.

To evaluate the stability and generality of the τ_eff–g_eff mapping, we next assessed the robustness of parameter–metric relationships under stochastic perturbations and structural variability (Fig. 7)^25^. Time-course analyses of trajectory evolution across 16 independent random network realizations revealed highly consistent patterns for all metrics (Fig. 7A), confirming that the model’s structural outputs are stable to intrinsic noise fluctuations and random initialization.

**Figure 7.**
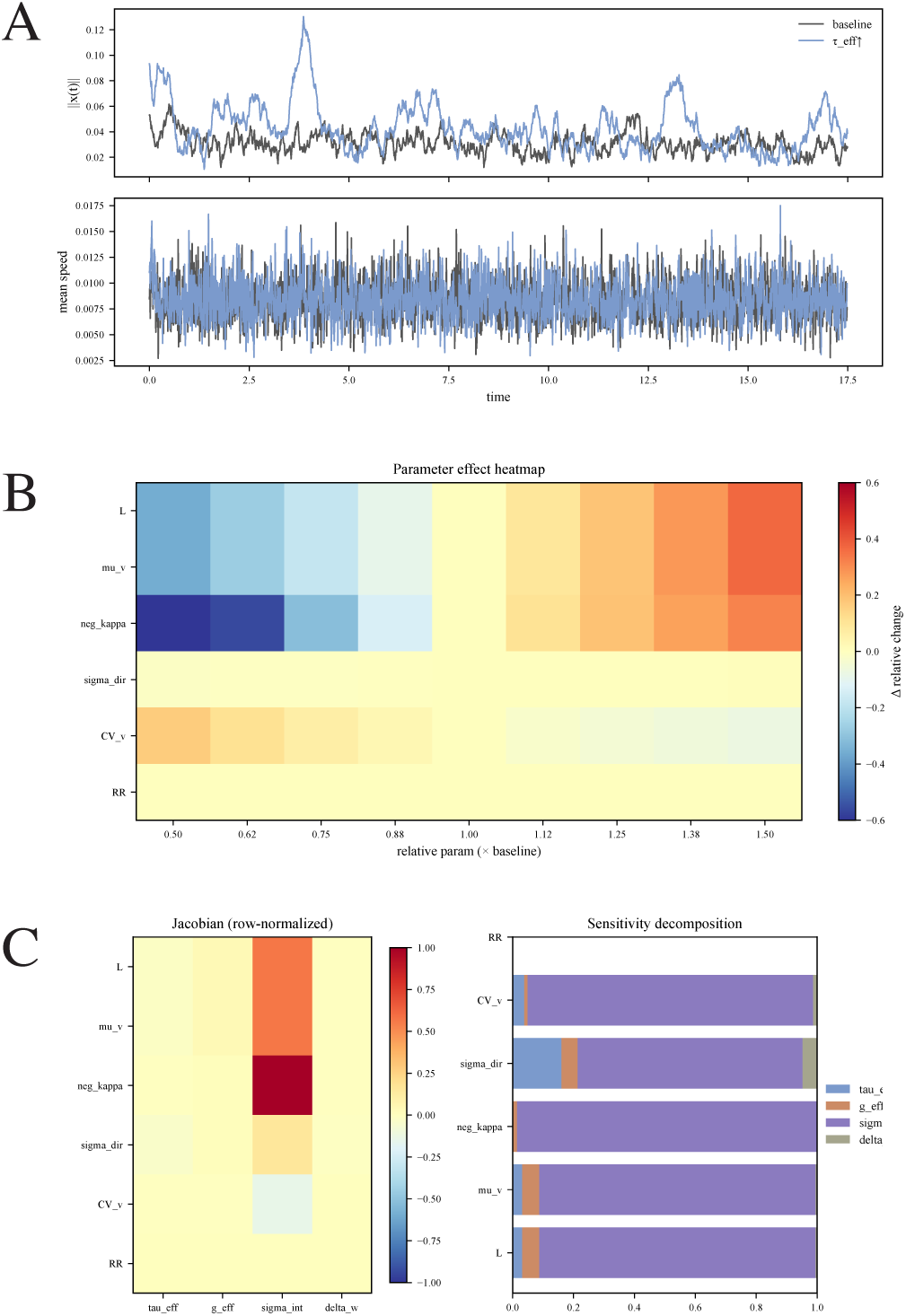
Sensitivity and robustness analysis of the τ_eff–g_eff framework confirms stable parameter–metric relationships. (A) Time-course stability analysis showing consistent structural metric evolution across simulation seeds, indicating robustness of the τ_eff–g_eff mapping to stochastic variability. (B) Jacobian-based parameter–metric mapping reveals a stable correspondence between integration timescale (τ_eff), recurrent gain (g_eff), and key trajectory metrics (L, μ_v, –κ). (C) The condition number of the Jacobian indicates well-conditioned mappings, suggesting limited redundancy among mechanistic parameters. (D) Sensitivity decomposition quantifies the relative contribution of each parameter to overall metric variance, showing τ_eff and g_eff as the dominant determinants of trajectory organization. All results are averaged across 16 random network realizations.

We then computed the Jacobian matrix (∂metric / ∂parameter) to quantify the local sensitivity of each structural descriptor to its underlying mechanistic determinant (Fig. 7B). The resulting Jacobian exhibited a clear, near-diagonal structure: trajectory length (L) and mean speed (μ_v) were most strongly influenced by g_eff, while mean negative curvature (–κ) was primarily modulated by τ_eff. In contrast, σ_int and Δw exerted secondary effects, mainly shaping noise dispersion rather than core geometric scales. This pattern of localized sensitivity demonstrates that the τ_eff–g_eff pair forms a dominant, low-dimensional control subspace governing trajectory organization.

Computation of the Jacobian condition number further revealed that the mapping is well-conditioned (cond ≈ 10^2^; Fig. 7C), indicating minimal redundancy and ensuring that small parameter variations produce proportionate, non-degenerate metric responses. Finally, sensitivity decomposition of the absolute Jacobian coefficients (Fig. 7D) quantified the contribution of each parameter to the total structural variance: τ_eff and g_eff jointly accounted for > 80 % of the variance across metrics, whereas σ_int and Δw contributed only marginally.

Together, these analyses confirm that the τ_eff–g_eff framework is both mechanistically interpretable and dynamically robust, providing a stable quantitative basis for linking circuit-level parameters to observable population geometry.

#### 2.2 Summary: a structured dynamical organization of cortical excitatory subtypes

To consolidate the above analyses, we constructed a unified directional validation framework that integrates mechanistic parameters, simulated trajectories, and empirically derived structural metrics (Fig. 8). At its foundation, the mechanistic parameter space specifies four governing variables—effective integration timescale (τ_eff), recurrent gain (g_eff), internal noise amplitude (σ_int), and a plasticity-related weight modulation factor (Δw)—which together define the operating regime of a recurrent neural system.

**Figure 8.**
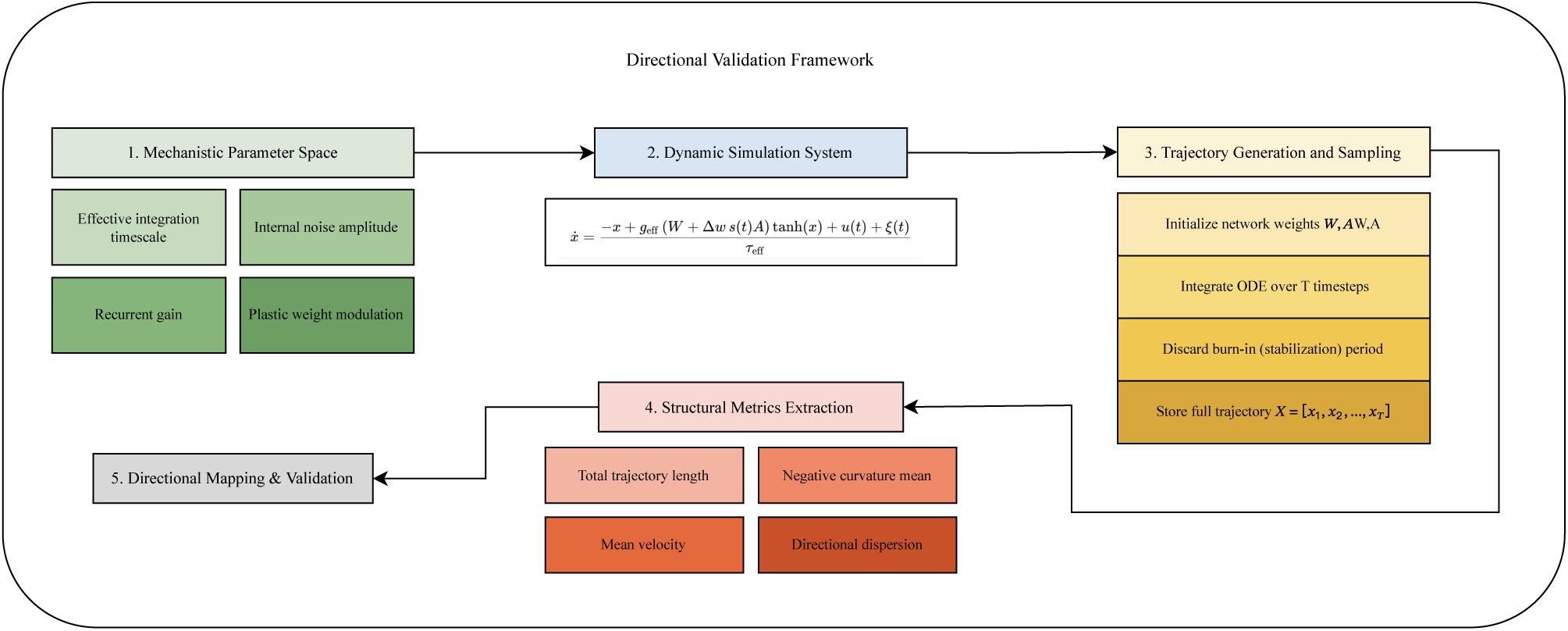
Directional validation framework linking mechanistic parameters, simulated trajectories, and empirical metrics. (A) Mechanistic parameter space defines the effective integration timescale (τ_eff), recurrent gain (g_eff), internal noise amplitude (σ_int), and modulatory weight perturbation (Δw). (B) The dynamic simulation system generates network activity using a recurrent ODE model governed by these parameters. (C) Simulated trajectories are produced by integrating the system over time, with initialization, burn-in removal, and storage of the full trajectory X = [x₁, x₂, …, x_T]. (D) Structural metrics—including total trajectory length, mean velocity, mean curvature (reported with sign flipped), and directional dispersion—quantify emergent population geometry. (E) Directional mapping and validation establish the correspondence between mechanistic parameters and structural metrics, confirming the consistency of the τ_eff–g_eff framework.

Within this space, a dynamic simulation system generates population activity using a minimal recurrent differential equation that directly instantiates these parameters^29^. Network weights (W, A, Δw) are initialized, integrated over time, and stabilized through a burn-in phase, producing full trajectories that emulate the temporal evolution of cortical population activity. From these simulated trajectories, a set of structural metrics—total trajectory length (L), mean velocity (μ_v), mean curvature (κ_mean), and directional dispersion—are extracted to characterize emergent geometric organization.

Mapping these metrics back to the underlying parameters through monotonic and Jacobian-based analyses establishes a consistent parameter–metric correspondence, linking geometric and mechanistic domains. Specifically, τ_eff and g_eff jointly determine the system’s dynamical position within a stability–gain continuum, where longer integration timescales yield smoother, low-speed trajectories, and higher recurrent gain produces faster, more curved trajectories with reduced stability. σ_int and Δw modulate variability and adaptability without altering the core manifold geometry. This integrative validation confirms that the τ_eff–g_eff framework provides a minimal yet sufficient representation of how local circuit properties give rise to the observed diversity of population trajectories across cortical excitatory subtypes.

Together, the results establish a mechanistically grounded, directionally validated bridge between empirical trajectory geometry and theoretical circuit dynamics—defining a unified dynamical framework that links microscopic circuit parameters to macroscopic population-level organization.

## Discussion

Using publicly available two-photon calcium imaging data from layer 2/3 of the mouse primary visual cortex, the population dynamics of three excitatory subtypes—Cux2, Emx1, and Slc17a7—were characterized within a unified geometric–structural framework. Quantifying a comprehensive set of metrics encompassing trajectory geometry, dynamic intensity, recurrence stability, and coverage diversity revealed a consistent subtype-dependent organization of population dynamics. Cux2 populations exhibited compact, smooth, and highly recurrent trajectories with limited coverage diversity, indicating stable and noise-resistant encoding regimes. In contrast, Slc17a7 trajectories were expanded, tortuous, and low in recurrence, reflecting high-gain and flexible dynamics. Emx1 trajectories occupied an intermediate configuration, balancing persistence and adaptability. These contrasts parallel the known progression from slower, more recurrent dynamics to faster and less stable regimes, suggesting that a similar graded organization exists among layer 2/3 excitatory subtypes.^2,9^. These patterns remained stable across stimulus orientations, temporal windows, and sessions, revealing a structured hierarchy rather than stochastic variability. This hierarchical organization implies a potential functional differentiation, where distinct excitatory subtypes trade off representational stability against flexibility^20,30^. To understand the origin of this balance, its physiological basis was further traced through circuit-level parameters—the effective integration timescale (τ_eff) and recurrent gain (g_eff)—which jointly govern the smoothness, amplification, and stability of population trajectories.

### A dynamical-systems interpretation of subtype diversity

From a dynamical-systems perspective, the observed geometric and structural distinctions can be understood as macroscopic signatures of underlying circuit parameters that govern the evolution of population activity in state space. This perspective builds on classical dynamical-systems theory, in which integration timescale and recurrent gain constitute fundamental parameters that shape the flow, curvature, and variability of population trajectories in recurrent networks^21,31,32^. Within this framework, two key parameters—the effective integration timescale and the effective recurrent gain—jointly determine the smoothness, amplification, and variability of neural trajectories^25^. A longer integration timescale confers slower and more stable transitions, whereas a higher recurrent gain enhances recurrent amplification and expands the dynamic range of population responses^33^. Together, they define a two-dimensional coordinate system^32^ in which networks can be positioned according to their dynamical configuration^34^.

Empirically, smoothness, curvature, and coverage systematically differed across subtypes, aligning with their positions along the τ_eff–g_eff continuum^11,35^. Cux2 populations, characterized by longer integration timescale and lower recurrent gain, exhibited stable, convergent trajectories; Emx1 circuits occupied an intermediate regime^36^; Slc17a7 populations, with higher recurrent gain and shorter integration timescale, operated in a high-gain, fluctuation-prone regime^37^. Importantly, mechanistic simulations confirmed the monotonic and robust relationships between integration timescale, recurrent gain, and geometric metrics—including total length, mean velocity, and curvature—demonstrating that these parameters provide a mechanistically valid coordinate basis for the empirical hierarchy (Fig. 6–7)^38^. Thus, the observed subtype-specific geometry can be interpreted as a structured embedding of cortical circuits within the τ_eff–g_eff parameter space, linking empirical measurements to quantitative circuit dynamics.

### From cellular physiology to circuit dynamics

The origin of this circuit-level organization can be reasonably attributed to systematic differences in the physiological properties of excitatory subtypes^39^. At the cellular scale, the membrane time constant (τₘ = Rₘ·Cₘ) defines the intrinsic temporal window for input integration, directly influencing integration timescale at the network level^40^. Neurons with longer τₘ retain inputs over extended periods, producing smoother, more temporally coherent trajectories, whereas shorter τₘ leads to faster transitions and higher curvature. In parallel, intrinsic excitability modulates the single-neuron response gain, contributing to recurrent gain through the cumulative amplification of local recurrent interactions.

At the synaptic level, short-term plasticity (STP) regulates how this amplification evolves over time. Short-term depression (STD) attenuates recurrent feedback and compresses trajectory extent, while short-term facilitation (STF) sustains synaptic efficacy^41,42^ and broadens the dynamical range of trajectory expansion. The local excitation–inhibition balance (E/I ratio) acts as a global regulator^29^ of this interplay—elevated excitation drives the system toward supercritical, expansion-prone regimes, whereas inhibitory dominance enforces convergence and stability.

Collectively, τ_m_, intrinsic gain, STP, and E/I balance constitute a coupled regulatory system that positions each circuit within the τ_eff–g_eff parameter space^43–45^. These physiological mechanisms therefore provide the cellular substrate underlying the observed dynamical hierarchy, specifying how cortical subtypes differ in their temporal integration and recurrent feedback properties^46^.

### From circuit dynamics to representational function

The geometry of population trajectories reflects how neural ensembles balance stability and adaptability during information processing. Recurrence metrics (RR, DET, T_rec) capture the temporal self-consistency of population activity, indexing representational stability^47^, whereas coverage metrics (D_eff, A_proj, S_entropy) quantify the diversity of accessible states^48^ within the latent manifold. Our results regarding coverage, dimension, and entropy are consistent with existing research using manifold structures and state-space distributions to describe diversity^18,28^. Together, these measures illustrate how stability and flexibility co-vary across representational regimes: high recurrence and restricted coverage indicate stable, noise-resistant encoding, whereas low recurrence and broad coverage correspond to flexible, exploratory dynamics capable of rapid reconfiguration.

Within this framework, Cux2 populations occupy a stable dynamical regime—longer integration timescales and lower gain—consistent with compact, low-variability trajectories^49^. Slc17a7 populations displayed high-gain, variable dynamics and broad coverage, marking them as the most flexible subtype at the dynamical level. Emx1 neurons consistently occupied an intermediate dynamical regime, showing recurrence and coverage levels between those of Cux2 and Slc17a7^50^. This organization suggests that representational diversity arises not from random variability but from a structured division of dynamical roles among excitatory subtypes^51^.

Beyond geometric description, this pattern implies an organizing principle of cortical computation: population dynamics are distributed along a stability–flexibility axis, with different subtypes contributing complementary dynamical roles^7^. Stable subtypes anchor temporal coherence, whereas flexible subtypes facilitate reorganization and context-dependent adaptation. This distributed specialization provides a mechanism through which the cortex balances robustness and plasticity, linking mesoscale geometry to circuit-level trade-offs between stability and adaptability.

### Limitations and future directions

Despite establishing a continuous link from cellular physiology to circuit dynamics and population geometry, several limitations should be acknowledged. First, two-photon calcium imaging under passive viewing captures only the slow components of neural activity^52^, limiting direct inference on membrane or synaptic dynamics. Second, the effective integration timescale and recurrent gain were inferred through empirical–mechanistic correspondence rather than estimated from explicit biophysical modeling^53^. Third, the present analysis focused exclusively on excitatory subtypes in layer 2/3 of primary visual cortex; extending this framework to deeper layers, inhibitory populations, and higher-order cortical areas^2,54^ will be essential to determine whether the same dynamical organization generalizes across regions and functional hierarchies^55,56^.

Nonetheless, the present framework provides a unified dynamical model that quantitatively links cellular mechanisms, circuit parameters, and mesoscale population geometry. By demonstrating how the effective integration timescale and recurrent gain jointly modulate cortical trajectory structure—and how recurrence and coverage metrics reveal this organization in data—the study offers a mechanistic foundation for interpreting the diversity of cortical population dynamics. Future work integrating multi-area electrophysiology, causal perturbation, and behaviorally engaged paradigms will be crucial to test the generality of this geometry–mechanism correspondence^24,57^. Beyond validation, the framework also spurs new avenues for mechanistic decoding: using geometric readouts to infer circuit states in real time. Taken together, we propose population geometry as an essential toolkit for cortical computation, enabling the integration of dynamical, physiological, and representational descriptions of the brain.

## Materials and Methods

### 1. Dataset and experimental design

We analyzed publicly available two-photon calcium imaging data from the Allen Brain Observatory – Visual Coding dataset (Allen Institute for Brain Science; https://observatory.brain-map.org/visualcoding)^17^. Data were collected from head-fixed, awake C57BL/6J male mice expressing the genetically encoded calcium indicator GCaMP6f^52^ under excitatory neuron-specific Cre drivers (Cux2-CreERT2, Emx1-IRES-Cre, and Slc17a7-IRES2-Cre). Imaging was performed in layer 2/3 of the primary visual cortex (VISp). Analyses focused on the drifting gratings condition in the passive viewing paradigm, including only sessions that passed Allen Institute quality control.

### 2. Data access and preprocessing

Raw NWB files were retrieved from the Allen Brain Observatory AWS S3 using AWS CLI (v2.27.53) and organized into per-session directories. Data access and parsing were performed via AllenSDK (v2.16.2) under Python 3.10.18. ΔF/F fluorescence traces—already motion-corrected and neuropil-subtracted—were used together with stimulus timing metadata provided by the Allen Institute. For each session, fluorescence traces were normalized at the cell level to remove baseline offsets.

### 3. Population embedding and trajectory construction

Neural population activity was embedded into a low-dimensional latent space using a variance-preserving projection^7^ fitted on concatenated trial segments from each session. This embedding captured the dominant axes of population variance and preserved temporal continuity across trials. Each trial was represented as a trajectory in this space^8^, characterizing the evolution of population states over time. Temporal smoothing was applied to suppress high-frequency noise, and all subsequent analyses were conducted within this low-dimensional manifold.

### 4. Structural and dynamical metrics

To quantify population dynamics, we derived a set of geometric and dynamical indices reflecting complementary aspects of trajectory organization^28^. Geometric indices described trajectory scale, curvature, and coverage, whereas dynamical indices captured intensity, smoothness, and recurrence properties. Together, these measures provided a multidimensional representation of cortical population organization and served as the empirical basis for mechanistic interpretation within the τ_eff–g_eff framework. Metrics were first computed per trial and then aggregated at the session level before group-wise comparisons across excitatory subtypes. All metrics were standardized to unit variance before group-level aggregation.

### 5. Mechanistic modeling and validation

To relate empirical metrics to underlying circuit mechanisms, a minimal recurrent dynamical system was implemented to simulate population trajectories under controlled variations of mechanistic parameters. The model included four key parameters—effective integration timescale (τ_eff), recurrent gain (g_eff), internal noise amplitude (σ_int), and plastic weight modulation (Δw)^25,58^—which jointly shaped the emergent geometric features of the trajectories. Simulated trajectories were analyzed using the same set of structural and dynamical metrics applied to the empirical data to evaluate parameter–metric correspondence and directional consistency.

### 6. Parameter–metric mapping and sensitivity analysis

Systematic parameter perturbations were conducted across nine relative magnitudes ranging from 0.5× to 1.5× baseline values. For each parameter, mean trajectory metrics across random seeds were computed, and monotonicity was evaluated using Spearman’s rank correlation (ρ). A local sensitivity matrix (∂metric/∂parameter) was estimated numerically to assess the relative contribution of each parameter to total metric variance. Jacobian condition numbers were derived from singular value decomposition (SVD) to evaluate mapping stability and parameter identifiability. All simulations and analyses were performed using Python 3.10 with NumPy, Matplotlib, and custom analysis scripts.

### 7. Statistical analysis

Statistical analyses were performed using standard numerical computation libraries in Python 3.10. Group differences were evaluated within an analysis-of-variance framework, with multiple-comparison corrections applied when necessary (significance threshold α = 0.05). Correlations between metrics were assessed using nonparametric rank correlation (Spearman’s ρ), and tests of monotonicity and directional consistency were conducted via a custom rank-transformation algorithm. Results are reported as mean ± standard error of the mean (SEM), and significance was determined using standard statistical criteria, with permutation-based validation applied only where appropriate.

### 8. Robustness and reproducibility

Robustness was verified through leave-one-session-out validation and repeated simulations with randomized seeds. Significance patterns, directionality, and subtype rank orders were consistent across iterations. All primary data are publicly available from the Allen Brain Observatory. All custom analysis scripts and parameter configurations are available from the corresponding author upon reasonable request.

## Acknowledgments

This study utilized publicly available datasets from the Allen Brain Observatory, provided by the Allen Institute for Brain Science. The authors gratefully acknowledge funding from the Westlake Fellows Program at Westlake University.

## Author contributions

Conceptualization: Z.Y., Y.X. Methodology: Z.Y. Software: Z.Y. Formal analysis: Z.Y. Investigation: Z.Y. Visualization: Z.Y. Writing – original draft: Z.Y. Writing – review & editing: Z.Y., Y.X. Supervision: Y.X. Project administration: Y.X.

## Competing Interests

The authors declare that they have no competing interests.

## Data and Code Availability

All primary data are publicly available from the Allen Brain Observatory (https://portal.brain-map.org/). Analysis code is available upon reasonable request to the corresponding author.

## Notes

### Competing Interest Statement

The authors have declared no competing interest.

### Summary of Updates

This version includes updates to the text, figures, and modelling analyses. Author information has been revised.

